# Trait-similarity and trait-hierarchy jointly determine co-occurrences of resident and invasive ant species

**DOI:** 10.1101/2020.02.05.935858

**Authors:** Mark K. L. Wong, Toby P. N. Tsang, Owen T. Lewis, Benoit Guénard

## Abstract

Interspecific competition, a dominant process structuring ecological communities, acts on species’ phenotypic differences. Species with similar traits should compete intensely (trait-similarity), while those with traits that confer competitive ability should outcompete others (trait-hierarchy). Either or both of these mechanisms may drive competitive exclusion within a community, but their relative importance and interacting effects are rarely studied. We show empirically that spatial associations (pairwise co-occurrences) between an invasive ant *Solenopsis invicta* and 28 other ant species across a relatively homogenous landscape are explained largely by an interaction of trait-similarity and trait-hierarchy in one morphological trait. We find that increasing trait-hierarchy leads to more negative associations; however these effects are counteracted when species are sufficiently dissimilar (by 37-95%) in their trait ranges. We also show that a model of species co-occurrences integrating trait-similarity and trait-hierarchy consolidates predictions of different theoretical assembly rules. This highlights the explanatory potential of the trait-based co-occurrence approach.

## INTRODUCTION

There is perhaps no ecological process that is at once as familiar and as enigmatic as interspecific competition, which can strongly mould the structure of ecological communities (Hutchinson, 1959). While patterns in biodiversity consistent with competitive interactions have been widely documented (Schoener, 1974; Calatayud et al., 2020), precisely how phenotypic differences between species determine the nature of competitive exclusion has remained highly contested.

Under classical niche theory (MacArthur & Levins, 1967), competitive exclusion leads to co-occurring species having dissimilar niches because species with similar niches compete more intensely. One proxy for the niche dissimilarity between two species is a non-directional or ‘absolute’ measure of their dissimilarity in trait space (Fig. 1A) (Carmona et al., 2019a). Accordingly, the *competition trait-similarity hypothesis* predicts that the likelihood of co-occurrence will always decrease with increasing overlap in trait space, such that co-occurring species display ‘overdispersion’: high absolute dissimilarity in trait space (Fig. 1A).

**Figure 1.**
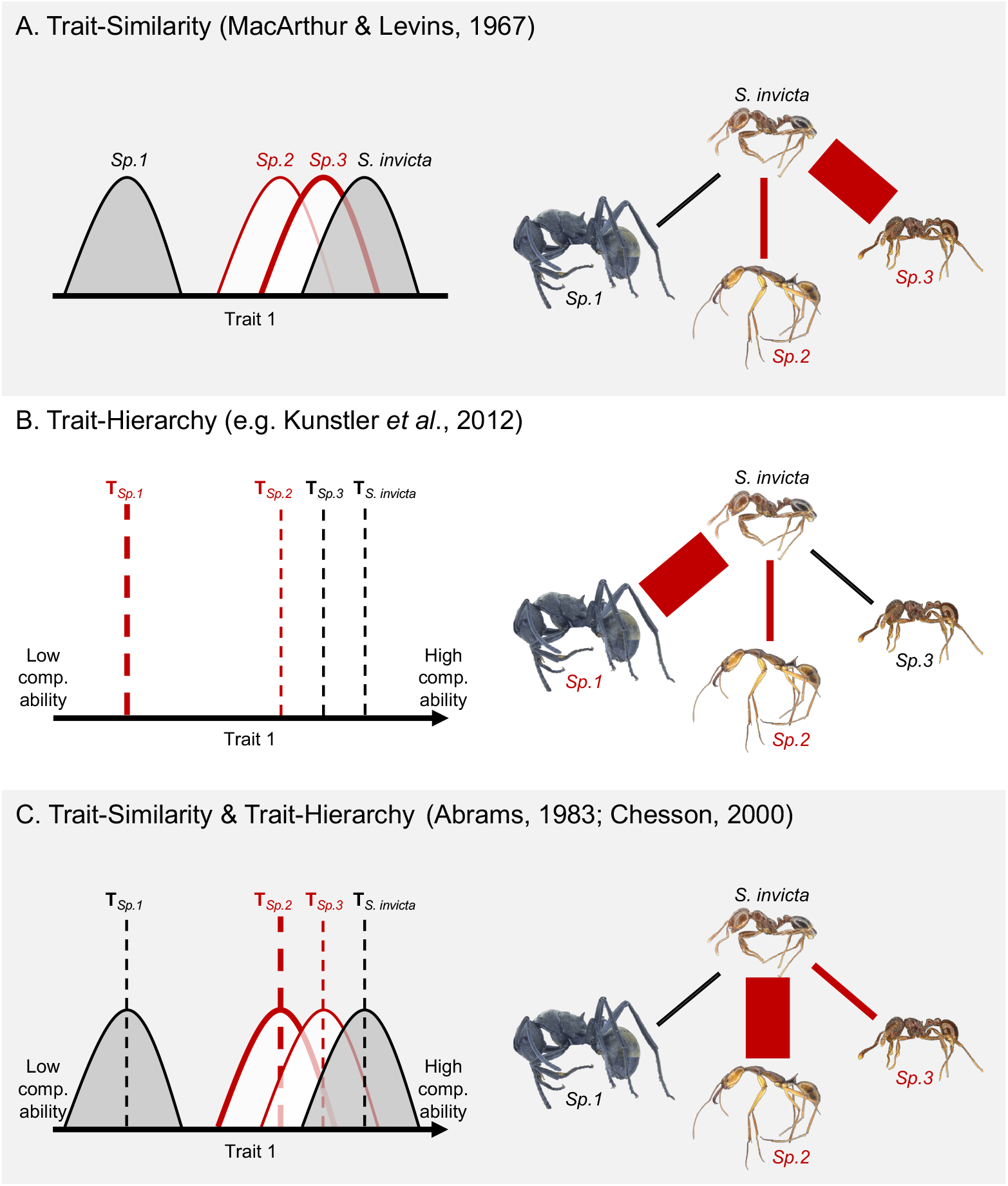
The trait-similarity and trait-hierarchy hypotheses of competition predict different outcomes for species co-occurrences separately and in combination. Panels show hypothetical relationships between three ant species and the invader *S. invicta* for one trait (left) and the corresponding pairwise co-occurrence relationships (right) as predicted under specific hypotheses. In each panel, species in red experience competitive exclusion and negative co-occurrence with *S. invicta* (i.e., they are not found in the same plots), with thicker lines indicating stronger relationships; species in black can co-occur with *S. invicta* in the same plots. **A**: If competitive exclusion is driven entirely by trait-similarity for all pairs of species (MacArthur & Levins, 1967), decreasing absolute dissimilarity (i.e. increasing overlap) between a species’ range of trait values and that of *S. invicta* increases the strength of the negative co-occurrence relationship, while increasing absolute dissimilarity (decreasing overlap) promotes co-occurrence. **B**: If competitive exclusion is driven only by trait-hierarchy (e.g., Kunstler et al., 2012) and species’ mean trait values (T) correspond to their competitive abilities along a directional axis, then a larger hierarchical difference (T1-T2) between a species and *S. invicta* increases the strength of the negative co-occurrence relationship, while a smaller hierarchical difference promotes co-occurrence. **C:** Trait-similarity and trait-hierarchy may jointly determine species co-occurrences because niche dissimilarities and competitive hierarchies interact to determine competitive outcomes across different species pairs (Abrams, 1983; Chesson, 2000). The likelihood of competitive exclusion (and strength of the negative co-occurrence relationship) between a species and *S. invicta* increases with increasing hierarchical difference in competitive ability; however, this competitive effect can also be counteracted and overcome by a large absolute dissimilarity in trait space, promoting co-occurrence.

In contrast, modern coexistence theory emphasizes that species’ niche dissimilarities are not the only factors determining competitive outcomes (Chesson, 2000). For instance, species can be organized along a competitive hierarchy where differences in competitive ability drive the exclusion of weaker competitors (Kunstler et al., 2012). Directional measures of trait differences, such as the ‘hierarchical difference’ in species’ mean trait values, provide a proxy for differences in competitive ability (Fig. 1B) (Kunstler et al., 2012). Contrary to the competition trait-similarity hypothesis, the *competition trait-hierarchy hypothesis* predicts that the likelihood of co-occurrence will decrease with increasing hierarchical difference (and dissimilarity), while decreasing hierarchical difference promotes ‘clustering’: the co-occurrence of similar species (Fig. 1B).

Despite a lasting focus on classical niche theory, empirical support for the competition trait-similarity hypothesis has been mixed (Mayfield & Levine, 2010). Some communities structured by competition show trait clustering consistent with the competition trait-hierarchy hypothesis (Herben & Goldberg, 2014). However, recent studies show that the outcomes of competition between plant species can be predicted by hierarchical differences in traits governing resource acquisition (e.g., leaf area for light interception, Kraft, Godoy & Levine, 2015; Kunstler et al., 2016; Perez-Ramos et al., 2019). The majority of trait-based studies applying the framework of modern coexistence theory have focused on plants and microbes, and have used experimentally-assembled communities (Grainger et al., 2019), which may not represent adequately the dynamics of natural communities (Carpenter, 1996). Most observational studies investigating the role of competition in structuring communities, however, measure only trait dissimilarities and test for overdispersion (Mittelbach & McGill, 2019). In this regard, the potential for species’ trait differences to reflect competitive ability differences may be underestimated.

Inferences of assembly processes from patterns in community structure are ubiquitous in the literature (Mittelbach & McGill, 2019). However, an inherent and questionable assumption of this approach is that all species within a community are subject to the same ‘dominant’ assembly process (Siepielski & McPeek, 2010). Rather than assuming that competition acts uniformly across all species at the community level, it can be informative to investigate whether and how competitive exclusion occurs for individual pairs of species. At this finer scale, competitive outcomes should be driven by an interaction between trait-similarity and trait-hierarchy (Chesson, 2000). That is, competitive exclusion will only occur for pairs of species which are insufficiently dissimilar in niches relative to their differences in competitive abilities (Fig. 1C; Mayfield & Levine, 2010). This interplay of trait-similarity and trait-hierarchy in determining competitive outcomes between species pairs is relatively unexplored. Nonetheless, it was anticipated by Abrams (1983): “What is needed instead is a broader definition of limiting similarity. The concept should be represented as a relationship between the difference in competitive ability and the maximum similarity that will permit coexistence. Such a relationship has the potential to be different for every different pair of species.”

Biological invasions, which often lead to intense competitive interactions, are choice settings for investigating competition (Shea & Chesson, 2002). For instance, many classical invasion hypotheses (empty niche, enemy escape, novel weapons etc.) essentially attribute invasion outcomes to niche dissimilarities and competitive ability differences between invader and native species (MacDougall et al., 2009). This framework of modern coexistence theory has been used to identify the trait values of exotic plant species which confer competitive advantages and facilitate invasion success (Gross et al., 2015) – but its potential to explain invasions in other taxa is untapped.

Ecological literature on the ants (Hymenoptera: Formicidae) is replete with studies identifying competition as a strong driver of community structure (Cerda, Arnan, & Retana, 2013) as well as reports of exotic species competitively excluding native ones (Holway, 1999). Many ant communities also show patterns of phylogenetic clustering in the presence of invasive ant species (Lessard et al., 2009), and theory suggests that such patterns may emerge if community assembly is driven by environmental filtering, or alternatively by competitive hierarchies. However, it is difficult to distinguish these two processes solely on the basis of phylogenetic relationships (Cadotte & Tucker, 2017). In most cases it is also hard to identify the species which compete most with invasive species, or which are most susceptible to displacement, especially when phylogenetic associations between invaders and resident species are ambiguous (Lessard et al., 2009). Such limitation in inferring contemporary ecological mechanisms from phylogenetic patterns of evolutionary history can be addressed with a focus on species’ traits, which govern their abiotic and biotic interactions in real time (Wong, Guénard & Lewis, 2019).

Here, we test trait-based hypotheses from classical niche theory and modern coexistence theory empirically. We focus on the invasion of the non-native Red Imported Fire Ant (*Solenopsis invicta*) in ant communities of wetland habitats in Hong Kong (reported in Wong, Guénard & Lewis, 2020). In these relatively homogenous landscapes, communities are more likely to be structured by competition as opposed to other mechanisms such as environmental filtering (Keddy, 1992). There is some disagreement as to whether *S. invicta* competes strongly with resident ant species during invasion. While some studies report competitive exclusion by *S. invicta* (Porter & Savignano, 1990; Gotelli & Arnett, 2000), others contend that altered abiotic conditions under anthropogenic disturbances – which happen to favour *S. invicta* – are directly responsible for the decline of resident species (King & Tschinkel, 2008). To this end, trait-based tests for theoretical mechanisms of competition in a system with low levels of environmental variation may clarify the interactions between *S. invicta* and other species.

We integrate trait-based and co-occurrence analyses to investigate whether trait-similarity and/or trait-hierarchy determine how *S. invicta* affect other ant species. There are two advantages to this approach. First, it allows for detecting potentially varying relationships at the fine ecological scales (species pairs) where competition unfolds (Abrams, 1983). Second, it allows for developing and testing more specific predictions about assembly processes than would be possible with standalone co-occurrence analyses (Veech, 2014). We first use a network of species’ spatial associations (co-occurrences) to quantify negative associations between *S. invicta* and other ants across multiple plots. Next, for distinct morphological traits that regulate ant physiology and behaviour, we use non-directional and directional measures of species’ trait differences as proxies for species’ niche dissimilarities (absolute dissimilarity) and competitive ability differences (hierarchical difference) respectively (after Kunstler et al., 2012; Carmona et al., 2019a). Integrating species’ trait differences and co-occurrences then allows us to test three hypotheses on the likelihood and nature of pairwise competitive exclusion between *S. invicta* and all resident species (Fig. 1).

If competitive exclusion is always driven by trait-similarity, absolute dissimilarity alone will determine co-occurrence relationships, with decreasing absolute dissimilarity leading to more negative co-occurrence (Fig. 1A). Alternatively, if competitive exclusion is always driven by trait-hierarchy, hierarchical difference alone will determine co-occurrence relationships, with larger hierarchical difference leading to more negative co-occurrence (Fig. 1B). Finally, if both mechanisms operate, we expect an interaction of absolute dissimilarity and hierarchical difference to determine co-occurrence relationships. Specifically, we expect absolute dissimilarity to modulate the effect of hierarchical difference, such that hierarchical difference determines co-occurrence relationships only if absolute dissimilarity is sufficiently low (Fig. 1C).

## MATERIAL AND METHODS

### Ant sampling and environmental variables

To maximise the likelihood of detecting community patterns reflecting biotic assembly processes such as interspecific competition (de Bello et al., 2012), we characterized ant communities at fine spatial scales in a relatively homogenous landscape (Wong et al., 2020). We selected two wetland reserves in Hong Kong – Lok Ma Chau (22.512°N, 114.063°E) and Mai Po (22.485°N, 114.036°E) – which have been conserved for >35 years, and which contain networks of exposed grass bunds (width ≤5 m) separating individual ponds. Most ant species present were native, but high densities of *S. invicta* were also recorded at multiple locations in pilot surveys conducted from 2015 to 2017. In 2018 we selected 61 plots from these locations, including 24 plots where *S. invicta* were present.

From April to September 2018, we sampled the local ant community at each plot in a 4 × 4 m quadrat, using six pitfall traps which were exposed for 48 hours (Wong et al., 2020). The maximum distance between any two traps in each plot was 5.65 m, a higher sampling density (i.e., traps / m^2^) than in previous studies characterising ant communities (Parr, 2008). We sampled at such fine spatial scales to enhance the detection of species’ occurrence patterns driven by biotic interactions, as most ant species in the region forage within 5 m of their nests (Eguchi, Bui & Yamane, 2004) and *S. invicta* forage within 4 m of their nests (Weeks, Wilson & Vinson, 2004). For the same reasons, a minimum distance of at least 20 m between individual plots facilitated independent observations.

All specimens were sorted into morphospecies and subsequently identified to species (Wong et al., 2020). We compiled a matrix of ant species’ occurrences (i.e., presence/absence data) across all 61 plots, and classified plots as either ‘*S. invicta*-present’ or ‘*S. invicta*-absent’ based on the presence of *S. invicta* workers in traps at each plot.

In addition to characterizing the ant community at each plot, we estimated the percentage of ground cover, and obtained data on the NDVI and mean annual temperature from local climate models at 30 m resolution (see Morgan & Guénard, 2019). We later used these data to check whether species’ preferences for particular physical properties influenced their co-occurrences (further below).

### Building co-occurrence networks

Co-occurrence networks document all pairwise co-occurrence relationships (i.e., network links) between species (i.e., network nodes) within a species pool. We used odds ratios to build the network (after Lane et al., 2014); this approach can incorporate signals of asymmetry in co-occurrence relationships (Araújo et al., 2011). We summarized the presence and absence of species pairs in 2*2 contingency tables and calculated the strength of co-occurrence relationships as their asymmetrical odds ratios (Lane et al., 2014). For example, given a species pair A & B, the odds ratio for indication of B by A (OR_AB_) measures how the probability of B’s presence at a plot changes under the presence of A in the same plot, and vice versa for OR_BA_:

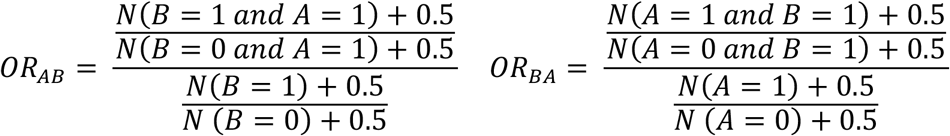

where N represents the number of plots. We applied Haldane’s correction and added 0.5 to all components to avoid odds ratios becoming infinity or undefined (Agresti, 2018). We further log-transformed the odds ratios in subsequent analyses such that they could be compared arithmetically (Agresti, 2018). All species were included in the analyses.

### Null models to assess co-occurrence relationships

To examine whether species were primarily associated with negative co-occurrence relationships within networks, we quantified their weighted degree – the sum of strengths (i.e., log-transformed odds ratios) of all co-occurrence relationships in the network. For each species, we only considered co-occurrence relationships which indicated how that species affected the odds ratios of other species being present in the same plots. For instance, the weighted degree of species A considered *OR*_*AB*_ but not *OR*_*BA*_.

Since any observed co-occurrence relationships could be driven by random associations (Gotelli, 2000), we used null models to compare their observed weighted degree to random expectation. Sampling plots were spatially distributed across two general localities – Lok Ma Chau and Mai Po (Wong et al., 2020) – and randomly shuffling species occurrences across the whole matrix could result in unrealistic null communities if the localities had different species pools. Thus, we randomly generated compositional data for each of the two localities, and then combined the two matrices to form one null matrix. To generate random matrices we used the fixed-fixed algorithm (“quasiswap” in R-package *vegan*), which is robust to Type-I errors and suitable for heterogenous compositional data (Gotelli, 2000).

We generated 1,000 null matrices, and calculated odds ratios to build null networks. We calculated the Standardized Effect Size for weighted degree (SES_WD_) (Gurevitch et al., 1992), defined as

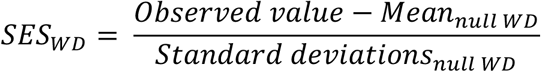

A species was primarily associated with negative co-occurrence relationships compared with random expectation if SES_WD_ was less than zero.

In order to test whether competition by trait-similarity and/or trait-hierarchy explained co-occurrences between *S. invicta* and resident ant species, we obtained the SES for all pairwise co-occurrence relationships involving *S. invicta* (SES_sinv_). Here we considered pairwise co-occurrence relationships which indicated how the presence of other species within plots were affected by the presence of *S. invicta*, but not vice versa. We obtained mean and standard deviations of odds ratios of each considered relationship from 1,000 null networks to calculate SES_sinv_ for each resident species. A negative SES_sinv_ value indicates that a species has a more negative co-occurrence relationship with *S. invicta* than expected by chance.

### Trait measurements and trait ranges of species

We assembled an individual-level trait dataset comprising data for seven morphological traits (body size, and six size-corrected traits: head width, eye width, mandible length, scape length, pronotum width, leg length). These traits regulate ant physiology and behaviour and are hypothesized to impact performance and fitness (see Table 1 in Wong et al., 2020). The dataset comprised trait measurements collected from at least 10 individual workers of every species (N=319 individual ants), including different subcastes (minor and major workers) of polymorphic species such as *S. invicta* (details in Wong et al., 2020). Body size was log-transformed to reduce right skewness. For each trait, we built species-level probability density functions (Carmona et al., 2019b) to calculate trait probability distributions (the curves in Fig. 1A). These distributions – or trait ranges – reflect the probabilities of observing different trait values *within* individual species; they were subsequently used to quantify absolute dissimilarities *between* species in trait (niche) space; see below.

**Table 1.** Multiple linear regression model for pronotum width. For this trait, a non-directional measure of niche dissimilarity (Absolute Dissimilarity, AD), a directional measure of competitive ability difference (Hierarchical Difference, HD), and their two-way interaction (AD*HD) determine pairwise co-occurrence relationships between the invader *S. invicta* and 28 ant species (Fig. 2B: SES_sinv_). Bold value indicates statistical significance (p<0.05). ‘JN intervals’ indicate the range of AD values at which the effects of HD are significant, as identified from the Johnson-Neyman procedure.

| Independent variable | $\beta$ | *P* | JN intervals |
| --- | --- | --- | --- |
| *AD* | -0.47 | 0.21 | <0.37; >0.95 |
| *HD* | 1.08 | 0.06 | - |
| *AD\*HD* | -1.47 | **0.002** | - |

R<sup>2</sup>=0.37

### Species’ trait differences, phylogenetic dissimilarity, and environmental preferences

For each of the seven traits, we quantified differences between *S. invicta* and each resident ant species with a non-directional measure of niche dissimilarity (Absolute Dissimilarity, AD), and a directional measure of competitive ability difference (Hierarchical Difference, HD). We measured AD as the proportion of a resident species’ trait probability density function which did not overlap with *S. invicta*’s trait probability density function (i.e., the proportion of trait space exclusive to the resident species’ trait range) (Carmona et al., 2019b). AD values range from 0 (when a resident species’ trait range is identical to that of *S. invicta*) to 1 (no overlap with trait range of *S. invicta*; e.g. *Sp. 1* in Fig. 1A). We measured HD as *T*_*Species*_ − *T*_S. invicta_, where *T* is the mean trait value for the given species (after Kunstler et al., 2012).

Additionally, to assess whether phylogenetic relationships or differing environmental preferences led to non-independence in co-occurrence relationships between *S. invicta* and each resident species, we quantified their phylogenetic dissimilarity (as pairwise distances between species in phylogenetic trees) as well as their dissimilarities in environmental preferences in terms of NDVI, ground cover and temperature (see Supporting Information).

### Statistical analyses

To determine whether pairwise co-occurrences between *S. invicta* and resident species were determined by trait-similarity, trait-hierarchy, or both mechanisms, we used multiple linear regression to test whether the SESsinv for each species-pair was best predicted by AD, HD, or an interaction of AD and HD. Our objective here was to use species’ trait differences to proxy their niche and competitive ability differences, rather than to understand the effect of different traits *per se*. Therefore, rather than using a full model, we built one model for each trait, with AD, HD and a two-way interaction term (AD*HD) as predictors. For the trait Mandible Length, AD and HD were highly correlated (Pearson’s *r* > 0.7), suggesting that their effects could not be separated (Dormann, 2013); thus, we excluded this trait from subsequent analyses.

For any trait models that detected a significant effect from the interaction of AD and HD, we used the Johnson-Neyman procedure (Johnson & Neyman, 1936) to calculate the ‘zone of significance’, that is, the range of values of AD at which HD influenced SES_sinv_ significantly (or vice versa). We controlled for false discovery rates using the procedure described in Esarey and Sumner (2017). We also checked whether the results of any models detecting significant effects were invariable to the use of different density-thresholds to classify *S. invicta*-present plots (see Supporting Information).

In addition to the individual trait models, we built separate models for phylogenetic dissimilarity and dissimilarities in species’ environmental preferences to determine whether these factors predicted co-occurrences between *S. invicta* and resident species. We built one model with phylogenetic dissimilarity as the sole predictor, and three additional models – each using environmental-preference dissimilarity in either NDVI, temperature or ground cover as a sole predictor. Environmental variables were also added to the trait and phylogenetic dissimilarity models as covariates if they were found to be significant.

Regression analyses were conducted in R (R Development Core Team, 2017). Before the analyses, we standardized the variables such that their relative importance could be assessed based on coefficient estimates (Schielzeth, 2010). We re-analysed our data with robust linear regressions to test whether our results were driven by statistical outliers.

## RESULTS

We recorded 29 ant species including *S. invicta* (Fig. 2), which occurred in 39% of the sampled plots. Within the co-occurrence network, *S. invicta* was the species most strongly characterized by negative co-occurrence relationships with other species (SES_all_=-3.62, Fig. 2A). Four other species were characterized by significant negative (SES_all_<-1.96) co-occurrence relationships, and two by significant positive (SES_all_>1.96) co-occurrence relationships (Fig. 2A). Of the 28 resident species, pairwise co-occurrences with *S. invicta* were positive (SES_sinv_>0) for nine species and negative (SES_sinv_<0) for 19 species (Fig. 2B). Of these, one positive and seven negative co-occurrence relationships were significant (Fig. 2B).

**Figure 2.**
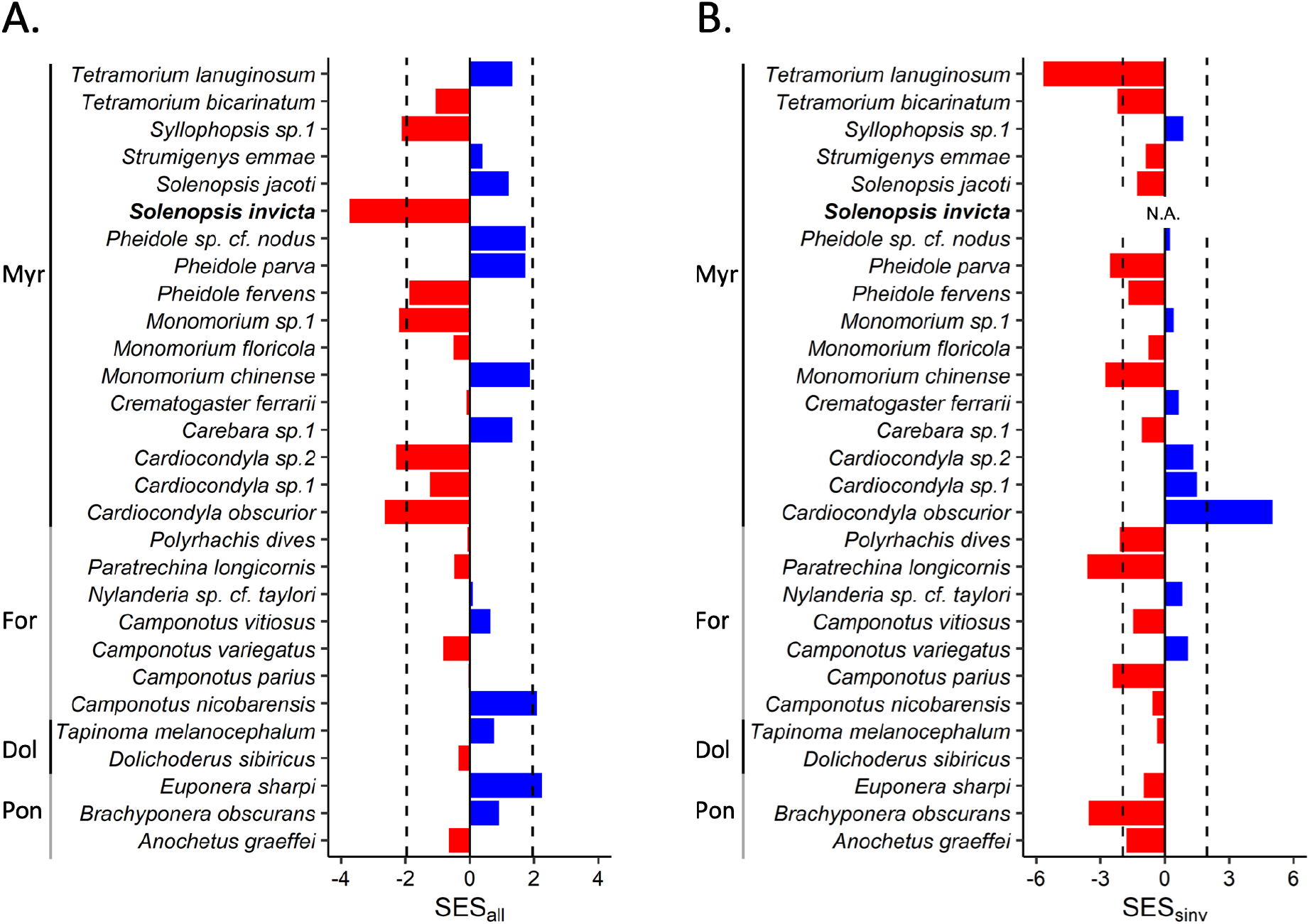
Of the 29 ant species sampled across 61 plots, the invasive ant *S. invicta* is most negatively associated with all other species. Plots show: (A) the degree to which each of the 29 species – including the invader *S. invicta* (in bold) – is characterised by positive (blue) or negative (red) associations within a co-occurrence network containing all species (SES_all_); (B) the degree to which each of the 28 resident species displays positive or negative associations with the invader *S. invicta* (SES_sinv_). Dashed lines indicate critical values for statistical significance of co-occurrence relationships (i.e., SES<-1.96 or >1.96). Ant species are grouped under four subfamilies: Myrmicinae (Myr), Formicinae (For), Dolichoderinae (Dol) and Ponerinae (Pon).

We found little evidence to suggest that either trait-similarity or trait-hierarchy solely determined species co-occurrences. On their own, both AD and HD were poor predictors of co-occurrence relationships between *S. invicta* and the 28 resident species (i.e., SES_sinv_) in separate models for six traits (Tables S1 & S3).

An interaction between niche dissimilarities and competitive ability differences best predicted co-occurrence relationships between *S. invicta* and the 28 resident species. Among different models for the six traits (Table S1), the most parsimonious model was that for pronotum width incorporating AD, HD and an interaction term (AD*HD), which explained 37% of the variation in SES_sinv_ (Table 1). Here, the interaction term (AD*HD) significantly explained co-occurrence relationships between *S. invicta* and the resident species (Table 1); removing the interaction term and only retaining the main effects of AD and HD significantly reduced model performance, as indicated by a Chi-square test (ΔAICc = 8.04, ΔAdjusted-R^2^ = 30.32, p < 0.001). A significant interaction between AD and HD was also consistently observed in all other models for pronotum width which used different density thresholds to classify *S. invicta*-present plots (Table S3).

In the model (Table 1; Fig. 3), the positive – and marginally significant – effect of HD on SES_sinv_ indicated that resident species with relatively wider or narrower pronotums than *S. invicta* tended to be positively or negatively associated with it, respectively. However, the significant negative effect of the interaction between AD and HD meant that the positive effect of HD on SES_sinv_ was reinforced when AD was low, and counteracted when AD was high.

**Figure 3.**
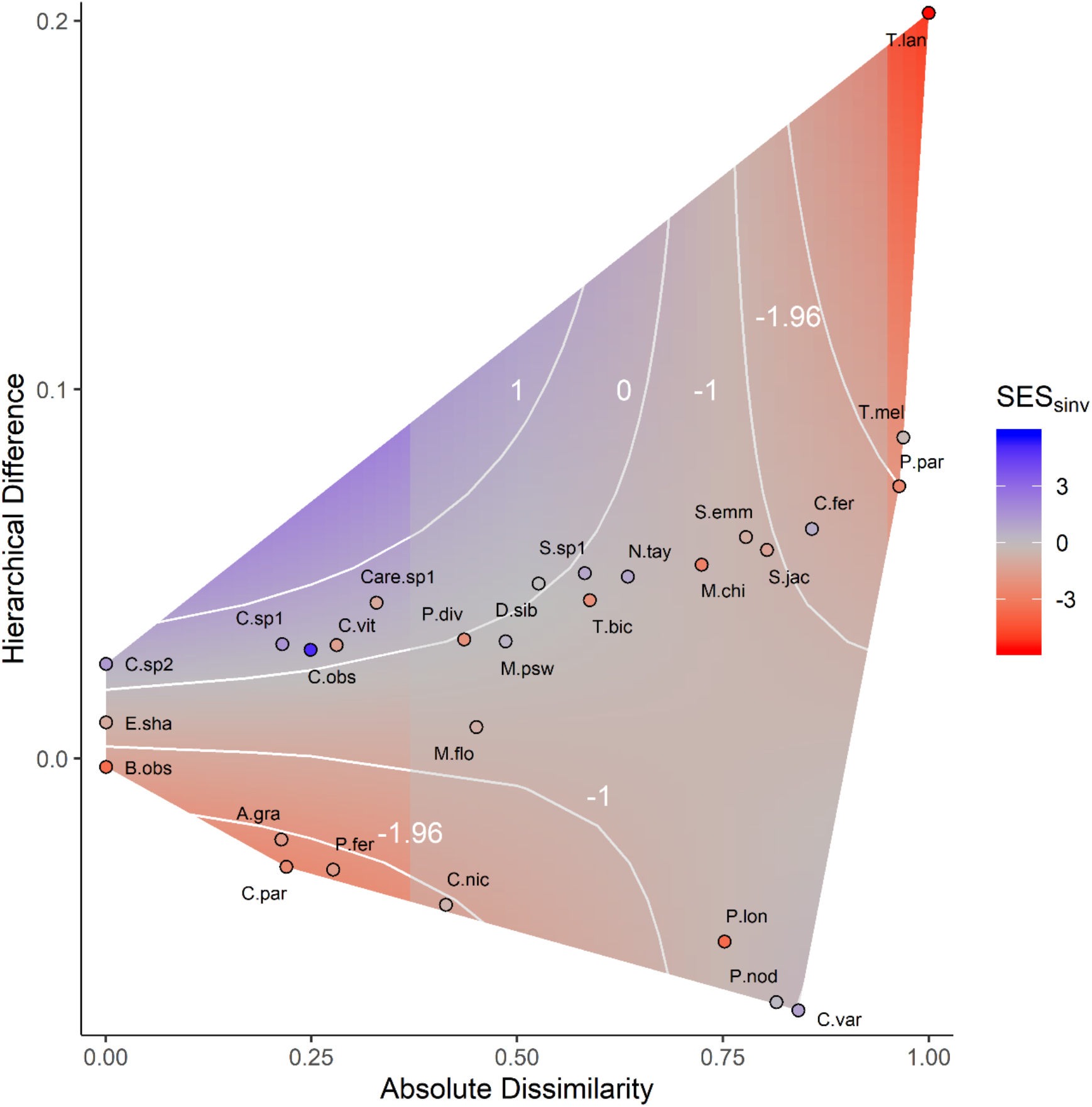
A response-surface showing how niche dissimilarity (Absolute Dissimilarity) modulates the effect of competitive ability difference (Hierarchical Difference) in determining resident ant species’ co-occurrence relationships with the invader *S. invicta*. The response-surface shows the predicted pairwise co-occurrence relationship between a given ant species and *S. invicta* (SES_sinv_) for the trait pronotum width, based on the multiple linear regression model in Table 1. Pairwise co-occurrence relationships (SES_sinv_) vary from negative (red) to positive (blue), with SES_sinv_<-1.96 and SES_sinv_>1.96 indicating significant negative or positive relationships respectively; contour lines illustrate how predicted SES_sinv_ changes across the response-surface. Coloured points on the response-surface show the observed SES_sinv_ for individual resident ant species (N=28) (full names of species shown in Fig. 2). On the x-axis, increasing values indicate decreasing overlap between a given species’ range of pronotum width values and that of *S. invicta.* On the y-axis, a positive or negative value indicates that a given species has a relatively wider or narrower pronotum than *S. invicta*, respectively. The masked area in the centre of the response-surface corresponds to the range of Absolute Dissimilarity (0.37-0.95) where the positive effect of Hierarchical Difference on SES_sinv_ is counteracted, as calculated from the Johnson-Neyman procedure.

Based on the model, we further estimated the magnitudes of niche dissimilarities (AD) between resident species and *S. invicta* at which competitive ability differences (HD) significantly influenced their associations. Applying the Johnson-Neyman procedure revealed that co-occurrence relationships between resident species and *S. invicta* were significantly affected by HD when AD<0.37 or AD>0.95. There were 10 species for which AD<0.37 and three species for which AD>0.95 in pronotum width with respect to *S. invicta* (Fig. 3).

In models based on other traits, main effects of AD and HD as well as their interacting effects were either non-significant or not consistently significant across different density thresholds of *S. invicta* presence (Tables S1 & S3). Phylogenetic and environmental-preference dissimilarities were also not significant predictors in any models (Table S2).

## DISCUSSION

We found that an interaction between trait-similarity and trait-hierarchy largely determined spatial associations (co-occurrences) between the invasive species *S. invicta* and 28 other ant species. These results suggest that trait-similarity and trait-hierarchy are interactive rather than discrete mechanisms of competitive exclusion, as predicted from theory (Chesson, 2000). We also found that a simple model of species co-occurrences, incorporating the interaction of trait-similarity and trait-hierarchy, broadly consolidated predictions of different theoretical rules of community assembly (discussed further below). Our study demonstrates that trait-based co-occurrence analyses uncover unique evidence that can help explain the outcomes of community assembly and biological invasions.

The overall pattern of pronounced negative co-occurrences between the abundant *S. invicta* and many other species (Fig. 2) strongly identifies *S. invicta* as an influential component of the network. Abundant species with many negative associations are often strong competitors (Calatayud et al., 2020). Previous studies (Gotelli & Arnett, 2000; LeBrun, Plowes & Gilbert, 2012) considered *S. invicta* to competitively exclude other ant species on the basis of negative co-occurrence patterns similar to those we observed. However, we appreciate that such patterns may also be generated by other ecological processes (Brazeau & Schamp, 2019). Thus, in order to strengthen inferences for particular assembly processes which could be at play, we explicitly scrutinized species’ co-occurrence relationships in light of their ecological differences (i.e., traits) within the context of classical and contemporary theories on interspecific competition (Fig. 1).

### Trait-similarity and trait-hierarchy jointly determine species’ co-occurrences

We found that no single mechanism of competitive exclusion (trait-similarity or trait-hierarchy) was sufficient to explain patterns of co-occurrences between *S. invicta* and the 28 resident ant species. However, incorporating the interactive effects of both mechanisms markedly improved explanatory power for a model based on the morphological trait, pronotum width (Table 1). Consistent with the basic principles of modern coexistence theory (Fig. 1), these results indicate that competitive outcomes among the ant species are unlikely to depend on niche dissimilarities alone, but on the relative magnitudes of these in relation to differences in their competitive abilities (Chesson, 2000). Competitive hierarchies in individual traits are known to structure some communities (e.g., plant height, Kunstler et al., 2016; Perez-Ramos et al., 2019) but are unexplored for most taxa. Our finding that differences in pronotum width significantly predict species’ associations (Table 1) highlights the potential importance of this frequently measured ‘functional’ trait (Parr et al., 2017) to competitive interactions among ant species. Given that the pronotums of ant workers contain the musculature powering load-bearing abilities (Keller et al., 2014), one testable hypothesis is that the relatively wider pronotums in *S. invicta* reflect a competitive advantage over other ant species (Fig. 3) through the more efficient capture, removal and transport of food resources. Notably, exploitative interspecific resource competition among ants is especially intense in less heterogenous habitats (Gibb, 2005), such as the one studied.

### Community assembly via trait differences: four rules

The trait model incorporating the interaction term (AD*HD) reconciled the varying co-occurrence patterns between *S. invicta* and individual ant species to the varying nature of each pair’s trait differences (i.e., in terms of trait-similarity and trait-hierarchy) (Fig. 3). Furthermore, the distinct ways by which species’ trait differences determine their co-occurrences as reflected in the model are strikingly consistent with predictions under different theoretical rules of community assembly. With reference to Fig. 3, our ecological interpretation of the model identifies four alternative rules which determine the pattern of co-occurrence between a given ant species and *S. invicta* across the landscape. Each rule is distinguished by the specific magnitudes of niche dissimilarities (AD) and competitive ability differences (HD) between paired species. The rules are: (I) competitive exclusion at HD<0 and AD<0.37, leading to negative co-occurrence; (II) approximate competitive equivalence and coexistence at HD>0 and AD<0.37, leading to non-negative co-occurrence; (III) sufficiently large niche dissimilarity and coexistence at AD=0.37–0.95, leading to non-negative co-occurrence; and (IV) environmental filtering at AD>0.95, leading to negative co-occurrence.

Rules I and II apply to species which are largely similar to *S. invicta* in niches and trait values (AD<0.37). Here the model predicts increasingly negative co-occurrences with increasingly negative HD (Fig. 3: left unmasked area: SES becomes negative as HD becomes negative). These results suggest that for ant species which have similar trait values to *S. invicta*, interspecific competition with *S. invicta* is likely to be intense, such that large differences in species’ competitive abilities drive exclusion, resulting in significant negative pairwise co-occurrences (e.g., Kunstler et al., 2012) (Rule I). However, for some species, small differences in competitive abilities with *S. invicta* (competitive equivalence) may facilitate coexistence with *S. invicta* in the fashion of neutral-like dynamics (Scheffer & van Nes, 2006) (Rule II). This is evident from the model, which predicts that the likelihood of co-occurrence for *S. invicta* and a similar species (AD<0.37) does not differ significantly from the null expectation (i.e., indicating coexistence is plausible) when HD becomes less negative (Fig. 3: left unmasked area: −1.96<SES_sinv_ <1.96).

In contrast to Rules I and II which apply to species sharing high niche similarity with *S. invicta* and competing intensely, Rule III applies to species which are largely dissimilar (AD=0.37– 0.95) from *S. invicta* in niches and trait values – to the extent that such niche dissimilarity may sufficiently mitigate any negative effects of competitive imbalances (e.g., individual plant traits in Perez-Ramos et al., 2019). Hence, for these species, differences in competitive abilities do not influence co-occurrences with *S. invicta* significantly (Fig. 3: masked area: SES_sinv_ does not significantly respond to HD). In addition, if niche dissimilarities are sufficiently large, coexistence is plausible, and the likelihood of these species occurring in the same plots as *S. invicta* generally does not differ significantly from null expectations (Fig. 3: masked area: - 1.96<SES_sinv_<1.96).

Rules I, II and III above concern interspecific competition, which we initially predicted to be an important driver of the ant species’ co-occurrences given the relatively homogeneous landscape. Less anticipated was an additional rule (IV), which likely relates to environmental factors. Rule IV applies to the minority of species which are most dissimilar from *S. invicta* in niches and trait values (Fig. 3: right unmasked area). For any species with such peak dissimilarity from *S. invicta* (AD>0.95), the model inherently predicts significant negative co-occurrence (SES_sinv_<-1.96) with *S. invicta* (Fig. 3). The extensive dissimilarities in trait values between these species and *S. invicta*, and the low likelihood of co-occurrence, may result from environmental filtering by unmeasured factors that vary across the plots (e.g., community-weighted means in ant species’ pronotum widths respond to gradients of soil fertility in Fichaux et al., 2019). If such trait-based environmental filtering occurs, directional differences in trait values could further reinforce its deterministic effects – this would explain the increasingly negative co-occurrence patterns observed with increasing HD (Fig. 3: right unmasked area).

In sum, different assembly rules collectively account for the co-occurrences of the invader *S. invicta* and the 28 resident ant species across the landscape, highlighting the multifaceted nature of community assembly. These findings broadly address the context-dependent nature of the impacts of *S. invicta* invasions on native ants in the collective literature (Porter & Savignano, 1990; Gotelli & Arnett, 2000; King & Tschinkel, 2008).

Abundant species, ranging from ants and beetles to trees and corals, often display negative and positive spatial associations with many other species (Calatayud et al., 2020). The ‘trait-based co-occurrence’ approach used in this study can provide insight into these ubiquitous patterns. Our parsimonious, single-trait model encompassing species’ trait differences (in terms of trait-similarity, trait-hierarchy, and their interacting effects) (Table 1; Fig. 3) reveals that an abundant species competes intensely with a subset of similar species, may coexist with species that are sufficiently different, and is further unlikely to co-occur with other species of different environmental requirements. This provides a realistic view of community assembly as a dynamic and multifaceted process acting varyingly on different pairs or sets of species (Abrams, 1983) – an alternative to the problematic notion of assembly as occurring via a static and discrete set of environmental and biotic ‘filters’ (Cadotte & Tucker, 2017).

Additional factors likely also influence co-occurrences between *S. invicta* and the 28 resident species, since the best individual trait model explained 37% of the variance (Table 1; Table S3). Also, if pronotum width was the only trait determining competitive outcomes, with wider pronotums indicating superior competitive abilities, we would expect ant species with AD<0.37 and HD>0 relative to *S. invicta* to exclude and show negative co-occurrences with *S. invicta*, instead of the non-negative co-occurrences predicted by the model (Fig. 3). In addition to the morphological traits measured in this study, other traits of ant species such as colony size or relative levels of intra- and interspecific aggression could potentially affect interspecific competition (Arnan, Cerdá & Retana, 2012). More broadly, the odds of competitive exclusion – and consequently precise patterns of species’ co-occurrences – are likely to depend on a net difference in competitive ability across multiple trait axes (Kraft et al., 2015). Thus, we envisage that the understanding of assembly processes such as interspecific competition in ecological communities can be enhanced by explicitly assessing trait-similarity, trait-hierarchy, and their interaction across diverse suites of morphological, physiological and behavioural traits. One could also extend the approaches used in this study into multi-dimensional trait space, where there is some evidence for the strong stabilizing effects of species’ dissimilarities (Kraft et al., 2015), but competitive hierarchies or the interacting effects of trait-similarity and trait-hierarchy are less explored.

### Trait-based co-occurrence: a framework for investigating community assembly

This study has shown that understanding of community assembly processes can be enhanced via a hypothesis-driven framework incorporating species’ trait differences and co-occurrence networks. Evidently, one advantage of such an approach is that it allows for the detection and consolidation of multiple assembly processes and their interactions, across fine ecological scales (species pairs and community subsets); whereas these processes may fail to be represented in coarser community-wide metrics such as functional or phylogenetic overdispersion and clustering (Mayfield & Levine, 2010). While experimental manipulations and mesocosm studies can be invaluable for understanding the precise mechanisms underlying community dynamics, their applicability decreases with increasing ecological, spatial and temporal scales (Levin, 1992). Data on species’ traits, abundances and distributions across multiple scales are increasingly collected and shared (Gallagher et al., 2019). For most species-rich ecological communities, we suggest that the trait-based co-occurrence approach represents an efficient and promising avenue for investigating assembly processes, and for identifying particular interactions, species and traits that are important determinants of community structure.

## Authorship

MKLW and TPNT conceived the study with inputs from BG and OTL. MKLW conducted fieldwork, collected trait data and built probability density functions. TPNT performed the statistical analyses. MKLW and TPNT wrote the first draft of the manuscript. All authors contributed significantly to revisions.

## Data accessibility

The authors confirm that, should the manuscript be accepted, the data supporting the results will be archived in the Dryad Digital Repository, and the data DOI will be included at the end of the article.

## AKNOWLEDGEMENTS

We thank Carlos Carmona and Christopher Terry for their comments on a previous version of the manuscript. This work was supported by a National Geographic Grant (60-16) and a University of Oxford Clarendon Scholarship to MKLW.

